# Phytoplankton functional types: a trait perspective

**DOI:** 10.1101/148312

**Authors:** Andrew J Irwin, Zoe V Finkel

## Abstract

Phytoplankton functional types are groupings of many species into a smaller number of types according to their ecological or biogeochemical role. Models describe phytoplankton functional types by a set of traits that determine their growth rates or fitness. Traits for functional types are often determined from observations on a small number of species under laboratory conditions. Functional types can be composed of a large number of species with very different trait values, so the representation of a type by an average trait value may not be appropriate. A potential solution is to estimate trait values from observations of the aggregate biomass of phytoplankton functional types in natural populations. We report on some recent efforts to extract trait values from time-series data using Bayesian statistical models and discuss some challenges of this approach.

## What are functional types?

Marine phytoplankton communities fuel the marine food web, producing 44-67 Gt of fixed carbon per year, which is approximately half of the total global net primary production (Field et al. 1998; Westberry et al. 2008). These communities contain many species each and in total there are tens of thousands of species of phytoplankton that inhabit the surface ocean (De Vargas et al. 2015; Guiry and Guiry 2015; Sournia et al. 1991). All phytoplankton species use chlorophyll or bacteriochlorophyll to harvest light as the energy source to fix organic carbon, but there is wide variation in virtually all their other traits. Phytoplankton species are found in eleven different phyla of the tree of life, including both prokaryotic and eukaryotic lineages. Several different endosymbiotic events gave rise to the major phytoplankton lineages within the eukaryotes (Falkowski et al. 2004; Katz et al. 2004).

Plant and phytoplankton functional types (or functional groups) are groupings of many species into a much smaller number of categories by function. While every species has its own set of unique traits, the functional type approach recognizes that we cannot analyze every species in the ocean and that some dimensions of variation will not matter for some research questions. Since species-level variation is often challenging to interpret, aggregation provided by functional types often facilitates identification of patterns at the level of communities and ecosystems. Phytoplankton communities are simplified into functional types for the design and interpretation of models that make predictions of phytoplankton biogeographic distribution, productivity, biogeochemical cycling and ecosystem function for the past, present and future (Anderson 2005; Gregg et al. 2003; Iglesias-Rodriguez et al. 2002; Le Quéré et al. 2005; Litchman et al. 2006). The simplification of phytoplankton communities from thousands of species to a handful of functional types greatly reduces both the computational effort required for modeling and the amount of data needed to constrain the parameters defining these models.

Phytoplankton functional types are often used to address questions about ocean biogeochemistry, for example, about the ability of the ocean to act as a carbon sink, the role of phytoplankton in ocean nitrogen fluxes, or CaCO_3_ and Si export (Jin et al. 2006; Le Quéré et al. 2005). This focus has led to the development of biogeochemically-defined phytoplankton functional types (Table 1). Some biogeochemical functions are phylogenetically conserved so that higher taxonomic classification can often be used as shorthand to define functional types for the phytoplankton. The ecologically dominant phytoplankton types are the silicifiers (diatoms), calcifiers (coccolithophores), N_2_-fixing and non-N_2_-fixing cyanobacteria. By definition all functional type groupings are simplifications; for example silicoflagellates, like diatoms, create silicified skeletons but are often omitted from the silicifier grouping because these taxa rarely dominate the biomass of modern phytoplankton communities. These functional type classifications arise from their biogeochemical roles, but also from contrasting biogeographies in the taxa and even analytical considerations such as the instrumentation needed to identify the species within these groups. Numerous other functional types are used less often, but with enough regularity that they are identified as phytoplankton functional types. These include the dinoflagellates (sometimes broken into sub-categories of mixotrophic or toxin-producing types), phytoflagellates (small-to-medium sized flagellated cells from several phyla which can be difficult to distinguish to the species level with a light microscope), picoeukaryotes (multi-phyletic photosynthetic eukaryotes less than 3 μm in diameter) and green (Chlorophyta) phytoplankton. There is some overlap among these groups; for example species from the Chlorophyta can be included in the picoeukaryotes (e.g., *Ostreococcus* sp.) and the phytoflagellates (e.g., *Halosphaera*). Although biogeochemically and taxonomically defined functional types dominate the literature, alternate definitions of phytoplankton functional types continue to be discussed (Flynn et al. 2015).

**Table 1.** Common phytoplankton functional types and taxonomic groupings. Functional types can be defined on the basis of biogeochemical or ecological function. Both approaches tend to identify types that are largely described by taxonomic groupings. For some types, one or a few taxa are commonly used as essentially equivalent to the entire type. Not all groupings are always used and a taxonomic group can fall into more than one biogeochemical or ecological functional type. Picoeukaryotes and Phytoflagellates are composed of multiple taxonomic groups.

| **Biogeochemical function** | **Ecological function** | **Taxonomic groupings** | **Exemplar species** |
| --- | --- | --- | --- |
| N<sub>2</sub>-fixers | Source of new nitrogen | Cyanobacteria | *Trichodesmium* |
| Silicifiers | Bloom formers, high export and trophic transfer efficiency | Diatoms | Many genera:<br>*Chaetoceros*,<br>*Skeletonema*,<br>*Thalassiosira* |
| Calcifiers | Bloom formers | Coccolithophorids | *Emiliana huxleyi* |
| DMS(P) producers | Climate regulation | Haptophytes | *E. huxleyi*,<br>*Phaeocystis* |
| Small cells | Nutrient recycling; low export and trophic transfer efficiency | Cyanobacteria, Picoeukaryotes, Phytoflagellates | *Prochlorococcus*,<br>*Synechococcus*;<br>*Ostreococcus* |
| Large cells | High export and trophic transfer efficiency | Diatoms, Dinoflagellates |  |

It is common to use one or two species as exemplars for each group, and these species tend to become functional synonyms for the groups (Table 1). For example, cyanobacteria are the most important contributors to nitrogen fixation in the ocean, and *Trichodesmium* is a common example (Hood et al. 2004; Lenes et al. 2001). Among the non-nitrogen-fixing species, the genera *Prochlorococcus* and *Synechococcus* dominate many oligotrophic regions (Chisholm 1992). The coccolithophore *Emiliania huxleyi* plays a large role in DMSP (dimethylsulfoniopropionate) and CaCO_3_ production. Several species of the Haptophyte *Phaeocystis* produce large gelatinous blooms that can impact food webs and can also be important producers of DMSP (Schoemann et al. 2005). DMSP is enzymatically converted to dimethylsufide (DMS), which has been proposed to alter cloud albedo and regulate climate (Charlson et al. 1987). Because diatoms are key contributors to carbon export and the dominant planktonic silicifiers, and many genera and species have been studied (some common genera include *Chaetoceros*, *Coscinodiscus*, *Pseudo-nitzschia*, *Rhizosolenia*, *Thalassiosira*, *Skeletonema*), this group is not commonly represented by a single species.

## The major functional traits

### What is a trait?

A phytoplankton functional trait is a fixed characteristic of a functional group that can be used to describe its growth rate or fitness. There is not yet a clear consensus on which traits are fundamental to understanding phytoplankton community dynamics or responses to environmental and climatic change and additional research will undoubtedly uncover new and more useful traits. We are just beginning to understand the variation and trade-offs in traits within and across functional types, the plasticity of traits in response to environmental conditions, and how traits can change in response to selection pressure.

### Types of traits

We find it useful to organize functional traits into conceptually similar groups: traits influencing overall metabolic rates, resource acquisition and requirements, and traits influencing loss rates such as sinking and susceptibility to grazing or attack by viruses and parasitoids. Some traits will influence or be correlated with other traits. Cell size acts as a master trait influencing metabolic rate, resource acquisition, sinking and susceptibility to grazing (Finkel et al. 2010a).

Maximum growth rate under resource-replete conditions is probably the most commonly used trait in modeling. Growth models are typically structured in terms of this trait and a wide range of factors that can reduce growth below this maximum. Generally the most limiting factor is of paramount importance for describing reductions in growth rate relative to the maximum. Various formulations of limitation are possible including a Liebig-style minimum relationship or co-limitation by two or more factors. Growth rate is easily measured for culturable organisms but is rarely measured under the full range of environmental conditions that can influence the maximum growth rate such as quality and quantity of irradiance, light-dark cycle, temperature, CO_2_ concentrations, salinity or different macro- and micronutrient concentrations and ratios in the growth media. It is generally assumed that acclimated exponential growth rates obtained for species grown in media with nutrients well in excess for growth under saturating but not super-saturating irradiances and under a light-dark cycle, and salinity and temperature typically experienced by the organism, will provide a good estimate of maximum growth rate. Other metabolic rates such as maximum photosynthetic rate, dark respiration and respiration as a function of growth rate (e.g., mol O_2_ per mol C) are often measured and can be used as traits (Geider 1992).

Several traits associated with nutrient acquisition have been identified. Nutrient acquisition traits are usually derived from a Michaelis-Menten or Monod parameterization of nutrient uptake as a function of nutrient concentration in the media (Fig. 1A, See Table 2 for a description of equations and symbols). The primary parameters are maximum uptake rate (*V*_max_) and the nutrient concentration where ½ *V*_max_ is achieved, termed the half-saturation constant (*K*_m_). These traits can be used to define derived traits such as nutrient affinity (*V*_max_/*K*_m_) and competitive ability, *R*\*, which is the lowest concentration at which a species has positive growth (*R*\* = *K*_m_ *d* / (*V*_max_ - *d*), where *d* is the loss rate through dilution or mortality). The Droop formulation (Fig. 1B) describes growth rate in terms of internal nutrient stores or cell quota (*Q*), a minimum cell quota (*Q*_min_) which is the smallest quota for positive growth, and a hypothetical maximum growth rate at infinite cell quota (μ’_max_). Maximum growth rate is some fraction of this hypothetical maximum growth rate. More elaborate formulations incorporate a maximum cellular quota (*Q*_max_) and allow uptake rates to vary with changes in nutrient quota or carbon to nitrogen ratios and storage (Aksnes and Cao 2011; Droop 1983; Flynn 2008; Grover 1991). Competitive ability, *R*\*, can be reformulated to include cell quota. Phytoplankton species require and acquire many nutrients and it is often useful to divide nutrients into different classes: macronutrients (various forms of fixed N and phosphate), silicic acid (mostly for diatoms), trace nutrients (Fe and many others), and organic nutrients (used by mixotrophs and heterotrophs). Monod and Droop models may not always be appropriate as there is evidence of linear or biphasic uptake kinetics for nitrate and some other nutrients for some phytoplankton species (Collos et al. 1992; Finkel et al. 2010b). Finally, carbon acquisition has its own complex set of traits because of the complexity of carbonate chemistry and carbon concentrating mechanisms (Raven et al. 2008; Reinfelder 2011).

**Figure 1.**
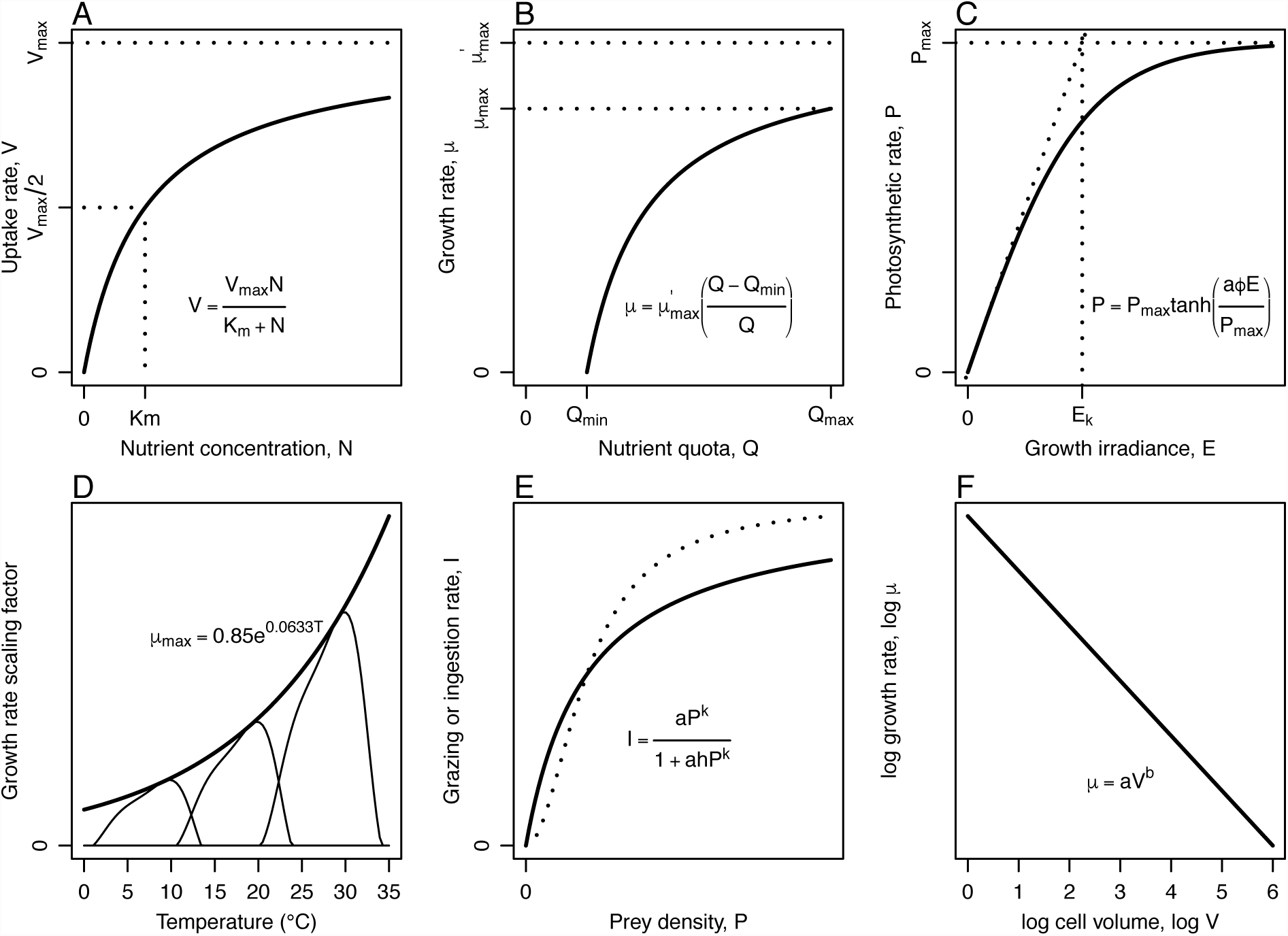
Common parameterizations of processes affecting growth and loss rates used to define phytoplankton functional traits. (A) Michaelis-Menten uptake kinetics for nutrients. (B) The Droop model for the effect of internal nutrient quota on growth rate. (C) Photosynthetic rate as a function of irradiance. (D) The effect of temperature on growth rate illustrated as the Eppley curve for maximum growth rate across phytoplankton species (bold line) and three species temperature responses for species with different temperature optima (thin lines). (E) Holling-type grazing rates as a function of prey density (type II, *k*=1, bold line; type III, *k*=2, dotted line). (F) Allometric scaling of biomass-normalized growth rate. Many other formulations of these functions have been used in the literature for each process. See Table 3 for a description of equations and symbols.

**Table 2.** **Examples of common parameters and equations describing growth and loss processes in phytoplankton. (See Fig. 1.)**

| Equation or Symbol | Description | Fig. 1 Panel |
| --- | --- | --- |
| $V = V_{\max} N / (K_m + N)$ | Michaelis-Menten uptake kinetics | A |
| $V$ | Uptake rate | A |
| $V_{\max}$ | Maximum uptake rate | A |
| $N$ | Concentration of nutrient in growth medium | A |
| $K_m$ | Half-saturation constant for nutrient uptake | A |
| $\mu = \mu'_{\max} (Q - Q_{\min}) / Q$ | Droop formulation of maximum growth rate based on internal nutrient quota | B |
| $\mu$ | Growth rate | B |
| $\mu'_{\max}$ | Theoretical maximum growth rate at infinite quota | B |
| $Q$ | Cell quota for resource that limits growth | B |
| $Q_{\min}$ | Minimum cell quota for positive growth rate | B |
| $P = P_{\max} \tanh(a\phi E / P_{\max})$ | Photosynthetic rate as a function of irradiance | C |
| $P$ | Photosynthetic rate | C |
| $P_{\max}$ | Maximum photosynthetic rate at saturating irradiance | C |
| $a$ | Absorption coefficient | C |
| $\phi$ | Quantum yield of photosynthesis | C |
| $E$ | Growth irradiance | C |
| $\mu_{\max} = 0.85e^{0.0633T}$ | Eppley curve for maximum phytoplankton growth rate across many species | D |
| $\mu_{\max}$ | Maximum growth rate | D |
| $T$ | Temperature | D |
| $\mu = a^T e^{-b(T-T_0)^c - \tau_2} / \tau_1$ | Species-specific curve for growth rate as a function of temperature | D |
| $\tau_1 = 0.33, \tau_2 = 0.30, a=1.04, b=1/1000, c=4$ | Shape factors and constants | D |
| $T_0$ | Reference temperature characterizing species-specific optimum temperature for growth | D |
| $I = aP^k / (1 + ahP^k)$ | Ingestion rate of phytoplankton (prey) by predators as a function of prey density (Holling type 2, 3 functional response) | E |
| $I$ | Ingestion rate | E |
| $a$ | Attack rate | E |
| $P$ | Prey number density | E |
| $k$ | Shape parameter (Holling type, see Fig. 1 caption) | E |
| $h$ | Time for prey to process prey (handling time) | E |
| $\mu = aV^b$ | Allometric equation describing size-scaling of | F |
| $\log \mu = \log a + b \log V$ | metabolic rates as a function of cell volume (two equivalent formulations) | |
| $a$ | Scaling factor to set growth rate at a reference size ($V=1$) | F |
| $V$ | Cell volume | F |
| $b$ | Size-scaling exponent (slope on the line in panel E), commonly $-1/4$ for biomass-normalized rates and $3/4$ for unnormalized rates. See Sommer et al. (2016) for a recent review of variability in $b$. | F |

**Table 3.**
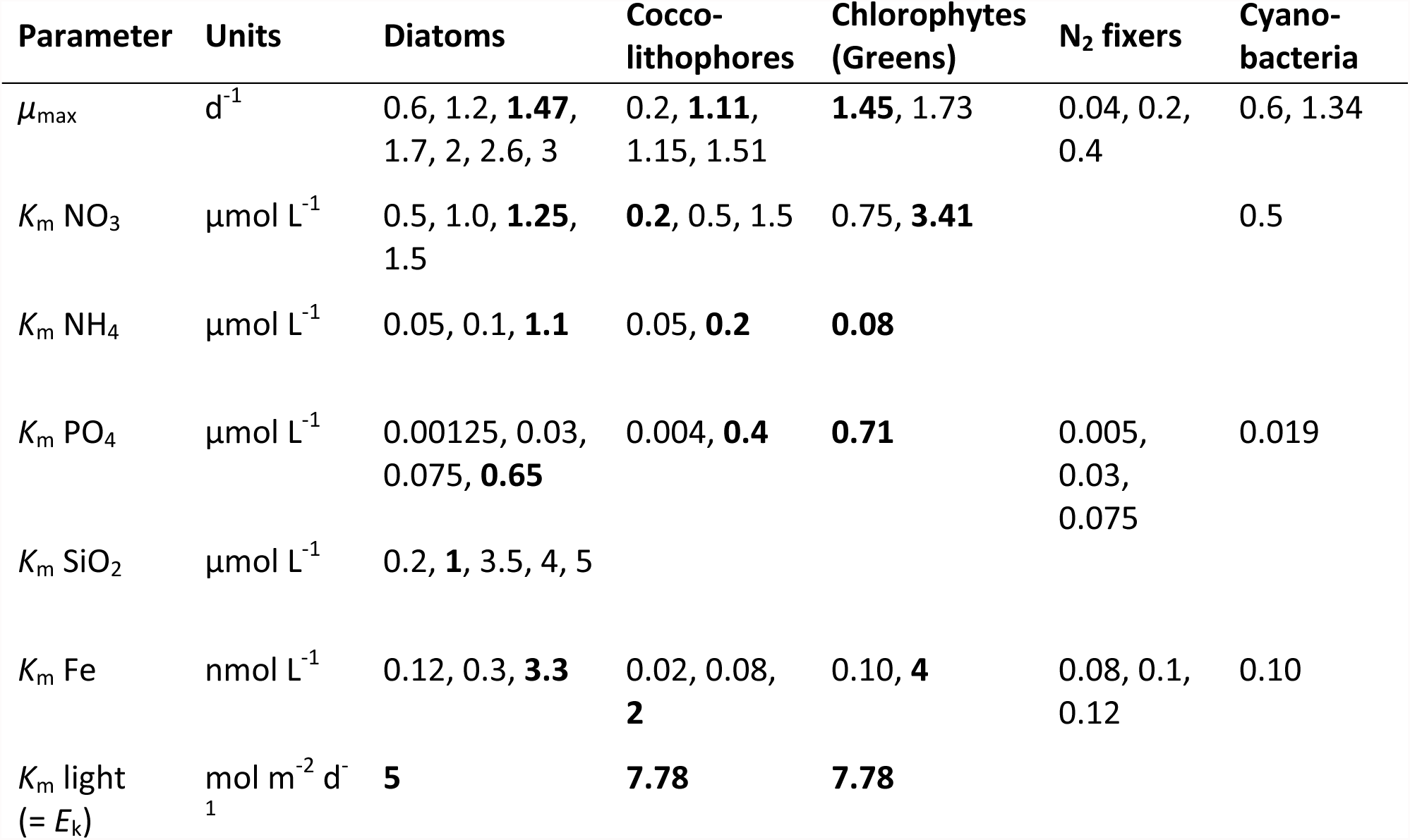
Summary of selected trait values from literature surveys and biogeochemical functional type models. Trait values from lab studies shown in bold are from Litchman et al. (2006). Other values (not in bold) are those used in phytoplankton functional type models developed by Dutkiewicz et al. (2009); Gregg et al. (2003); Le Quéré et al. (2005); Merico et al. (2004); Moore et al. (2002); Sarmiento et al. (2010). Maximum growth rates are reported at 30°C by Gregg et al. (2003) (the highest values, except for diatoms) and at 0°C by Le Quéré et al. (2005) (the lowest values).

| Parameter | Units | Diatoms | Cocco-lithophores | Chlorophytes (Greens) | N<sub>2</sub> fixers | Cyano-bacteria |
| --- | --- | --- | --- | --- | --- | --- |
| $\mu_{\max}$ | d<sup>-1</sup> | 0.6, 1.2, **1.47**, 1.7, 2, 2.6, 3 | 0.2, **1.11**, 1.15, 1.51 | **1.45**, 1.73 | 0.04, 0.2, 0.4 | 0.6, 1.34 |
| $K_m$ NO<sub>3</sub> | μmol L<sup>-1</sup> | 0.5, 1.0, **1.25**, 1.5 | **0.2**, 0.5, 1.5 | 0.75, **3.41** | | 0.5 |
| $K_m$ NH<sub>4</sub> | μmol L<sup>-1</sup> | 0.05, 0.1, **1.1** | 0.05, **0.2** | **0.08** | | |
| $K_m$ PO<sub>4</sub> | μmol L<sup>-1</sup> | 0.00125, 0.03, 0.075, **0.65** | 0.004, **0.4** | **0.71** | 0.005, 0.03, 0.075 | 0.019 |
| $K_m$ SiO<sub>2</sub> | μmol L<sup>-1</sup> | 0.2, **1**, 3.5, 4, 5 | | | | |
| $K_m$ Fe | nmol L<sup>-1</sup> | 0.12, 0.3, **3.3** | 0.02, 0.08, **2** | 0.10, **4** | 0.08, 0.1, 0.12 | 0.10 |
| $K_m$ light (= $E_k$) | mol m<sup>-2</sup> d<sup>-1</sup> | **5** | **7.78** | **7.78** | | |

Light harvesting by photosynthesis can be described by two traits analogous to the maximum uptake and half-saturation rate: the maximum photosynthetic rate often termed the photosynthetic capacity (*P*_max_) and saturation irradiance (*E*_k_), both derived from curves of photosynthetic rate as a function of irradiance (Geider and Osbourne 1992; Jassby and Platt 1976). Photosynthetic efficiency, *α* = *P*_max_ / *E*_k_, estimated from the curve when irradiance approaches zero, is commonly used as a trait (Fig. 1C). Fundamentally photosynthetic efficiency is the product of the maximum quantum yield of photosynthesis (moles carbon or oxygen produced per mole photons absorbed) and the light absorption coefficient, the amount of light absorbed by the cell often normalized per unit of chlorophyll-a. Theoretically under low light the maximum quantum yield is a constant (Φ), but the light absorption coefficient (*a*) varies across species and environmental conditions due to differences in pigment composition, concentration, and cell size (Finkel et al. 2004a; Finkel 2001; Kirk 1994). High or super-saturating irradiances above *E*_k_ reduce net photosynthesis and are commonly parameterized by a single value, β or *E*, representing the irradiance level at which light inhibition becomes significant (Platt et al. 1980). Many other traits, including the capacity for non-photochemical quenching, the effective cross-section of photosystem II, and the cross-section of photoinactivation of photosystem II can be measured and have been shown in individual studies to influence the susceptibility of different species and species of different size (Key et al. 2010; Lavaud et al. 2007; Six et al. 2007) to high light stress, but these and other traits related to strategies to deal with high light stress are rarely used in ecological or biogeochemical models (Raven 2011).

The rate of a chemical reaction is affected by temperature according to the Arrhenius law, which describes the temperature dependence of a chemical reaction as proportional to exp(-*E_a_*/*kT*), where *Ea* is the activation energy of the reaction, *k* is Boltzmann’s constant, and *T* is the absolute temperature in Kelvin. Despite the tremendous complexity of a living organism, the Arrhenius law can predict metabolic rate changes over broad ranges of temperatures. There is evidence that *E_a_* varies across broad taxonomic groupings (Gillooly et al. 2001). The Arrhenius law is sometimes approximated as an exponential function of temperature using the trait *Q*_10_, which describes the multiplicative change in metabolic rate with a 10°C change in temperature, or even as a linear response over a narrow temperature range (Montagnes et al. 2003). For phytoplankton, the maximum growth rate across species is often described as increasing exponentially with temperature, μ_max_ = 0.85 exp(0.0633 *T*) (Eppley 1972). Single species exhibit a gradual increase in growth rate with temperature followed by a rapid decline above a maximum temperature and each species is adapted to a specific temperature range which is narrower than the range described by Eppley’s curve (Fig. 1D) (Follows et al. 2007). High temperatures dramatically reduce rates and inactivate enzymes, so a maximum temperature, *T_max_*, can also be a useful trait to describe phytoplankton physiology. Other useful temperature-related traits, such as the minimum temperature for positive growth rate (*T*_min_), can be derived from observations of metabolic rate as a function of temperature (Boyd et al. 2013).

Traits affecting loss rates are more difficult to identify, but the usual collection includes the sinking rate, susceptibility to grazing and susceptibility to viral and parasitoid attack. Sinking rate, *V*, is predicted by Stokes’ law as the terminal velocity of a spherical particle with a very small Reynolds number and is often applicable to phytoplankton, where

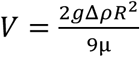

and *g* is acceleration due to gravity, Δρ is the density difference between the cell and surrounding medium, μ is the viscosity of the medium, and *R* is cell radius. Corrections for shape may be needed for cells that are not approximately spherical. A cell’s ability to regulate its density and thus buoyancy may be an important trait that strongly influences its sinking rate in practice (Bienfang et al. 1982; Miklasz and Denny 2010). Silica or calcium carbonate will affect the density of a cell and serves as ballast, increasing the sinking and export rates of cells relative to unballasted cells (Armstrong et al. 2002). A related but less well-studied trait that affects export efficiency is the tendency to form aggregates, which can be facilitated by exudation of DOC, the stickiness of cells, and interaction with grazers (Alldredge and Silver 1988).

Many traits influence grazing susceptibility including the presence of mineralized cell walls, quality as food, the size and shape of the cell, colonial growth forms, spines, and motility. With the exception of cell size, these traits are not usually incorporated into a mathematical model of grazing, but at most are aggregated into an edibility factor to moderate grazing rates or trophic transfer efficiency. Grazers usually prefer to consume prey cells that are about an order of magnitude smaller, although this ratio varies for different predators (Hansen et al. 1994), and this rule can be used to develop size-structured grazing models and help develop theory about the size structure of food webs (Loeuille and Loreau 2005). At this stage very little is known about the traits that are useful for modeling attack by viruses and parasitoids, and these factors are rarely included in models. Traits that govern loss rates for phytoplankton are usually parameterized very simply as a constant fraction of prey density, a density-dependent rate, or with two size-classes of grazers (Le Quéré et al. 2005; Moore et al. 2002) but there are notable exceptions to these very simple approaches (Armstrong 2003; Vallina et al. 2014). Two very common approaches for modeling grazing rate, or ingestion rate (*I*) of prey (*P*) by predators, are the Holling type II and type III functional responses which have attack rate (*a*) and prey handling time (*h*) as traits and the shape parameter k distinguishes the two responses (type II, *k*=1; type PII, *k*=2, Fig. 1E).

### Size as a master trait

Cell size is often referred to as a master trait because body size influences the physiology, ecology, and evolution of species (Finkel 2007). For a recent review of some of the recent literature see Sommer et al. (2016). Phytoplankton cell size (radius) varies over three orders of magnitude (Finkel et al 2010a). Phytoplankton cell size is mechanistically linked to all the physiological and ecological traits discussed above: maximum growth rate, nutrient acquisition and minimum and maximum cell quota, light absorption and susceptibility to high light stress, sinking rate and susceptibility to grazing and viral attack (Finkel et al. 2010a). The size-dependence of traits leads to clear biogeographic patterns in phytoplankton community size structure: small, predominantly picoplankton cyanobacteria, such as *Synechococcus* and *Prochlorococcus* spp., dominate the phytoplankton community biomass in oligotrophic gyres while diatoms often dominate in upwelling zones (Barton et al. 2013). Small phytoplankton tend to dominate under oligotrophic conditions due to a combination of their low nutrient requirements (small minimum and maximum nutrient quotas), low half saturation constants for nutrient (*K_m_*), and high maximum nutrient uptake and growth rates. High export rates of carbon to the deep sea and more efficient trophic transfer of carbon to higher trophic levels in the food web are more likely when standing stock biomass is high and large phytoplankton cells dominate (Finkel et al. 2010a; Laws et al. 2000). In the near future, communities may shift to smaller-sized phytoplankton due to changes in environmental conditions linked to climate change (Li 2002; Li et al. 2009; Morán et al 2010). Climate change has been associated with macroevolutionary shifts in the size of several plankton groups over the last 65 million years of Earth’s history: including the diatoms (Finkel et al. 2005), coccolithophores (Henderiks and Pagani 2008), dinoflagellates (Finkel et al. 2007), silicoflagellates (Van Tol et al. 2012) and foraminifera (Schmidt et al. 2004).

Most physiological rates, including maximum growth rate and maximum photosynthetic and uptake rates, and cellular components, including minimum and maximum quota, are power-law functions of cell size. For eukaryotic phytoplankton, maximum metabolic rates generally follow a ¾ size-scaling rule. When metabolic rates are normalized by cell volume, this exponent is generally −¼, meaning for example that growth rate (*μ*) is proportional to cell volume (*V*) to the −¼ power (Fig. 1F). This size allometry for metabolic rates was first documented for animals (Kleiber 1947) and is one of the most general laws in biology. There are numerous exceptions to the general size-scaling trend in phytoplankton, particularly when resources are limiting (Finkel et al. 2004b; Irwin et al. 2006; Mei et al. 2009; Mei et al. 2011). There is some evidence that the size scaling of metabolic rates as a function of organism mass (e.g., cell carbon content) is closer to 1 for single-celled organisms, such as the phytoplankton, and <$> for metazoans (Delong et al. 2010). Current research is taking a closer look at the mechanistic basis for size scaling of physiological processes in phytoplankton and other organisms to better understand the reasons for the scaling laws and the anomalies (Kempes et al. 2012; Sharpe et al. 2012).

Nutrient acquisition is perhaps the easiest process to analyze in terms of its size-dependence. Acquisition of resources, whether by active or passive uptake is across the cell wall, and the surface area of the cell is proportional to the square of the radius of the cell, while the volume is proportional to the cube of the radius. Nutrient porters are arrayed across the cell surface and so under limiting conditions, the amount of surface area dedicated to nutrient acquisition can become limiting, leading to insufficient capacity for nutrient uptake. Competition within the cell for surface area can lead to trade-offs for the acquisition of one nutrient compared to another, depending on the relative degree of limitation (Edwards et al. 2011). The half-saturation constant for nutrient uptake (*K*_m_) is also size-dependent, scaling proportionally with the radius of the cell, so that larger cells have larger half-saturation constants and thus are poorer competitors, in general, for dissolved resources when concentrations are low. This is partly a result of the rate of diffusive flux of resources across the boundary layer outside a cell, the size of which is proportional to cell radius. The quotient of these two quantities, nutrient affinity (*V_max_*/*K_m_*), is thus proportional to the cell radius as well, so that nutrient affinity increases with cell size. This size dependence also affects the acquisition of dissolved inorganic carbon. Although the active uptake of bicarbonate is not generally limiting in seawater, the rate of CO_2_ diffusion in and out of the cell is affected by cell size according to Fick’s first law: diffusive flux *F* = 4π *R D* Δ*C*, where *R* is radius, *D* is the corresponding diffusion constant, and Δ*C* is the concentration gradient between the bulk media and the surface of the cell (Reinfelder 2011).

Light acquisition, and thus photosynthetic rate when light is limiting, is affected by cell size, but the explanation is more complex than for nutrient acquisition because of photoacclimation mechanisms within the cell. Larger cells have, in general, lower intracellular concentrations of pigments compared to smaller cells because the path-length of light is longer across a larger cell (Finkel 2001). This means that the energy absorbed per pigment, when all other factors are equal, will be lower in a larger cell. This effect is referred to as the package effect, and it is the reason why intracellular pigment concentrations are generally inversely proportional to cell radius for light-limited cells (Finkel et al. 2004a). This size-dependence of light acquisition contributes to slower growth rates for larger, light-limited cells, compared to smaller cells. By contrast, pigment packaging confers an advantage to larger cells under high light because the same effect reduces the rate of photoinactivation of photosystem II (Key et al. 2010).

In addition to sinking, grazing rates and trophic transfer efficiency can be predicted approximately from cell size. Since size is a major factor influencing all these characteristics of phytoplankton, it can often be used as a proxy for a wide suite of traits (Irwin et al. 2006; Wirtz 2013). The ability to simplify model parameterizations is tremendously useful, but more work should be done to identify the most important failures of this approach to improve modeling efforts. Cell shape and chain and colony formation can be important traits affecting phytoplankton growth and loss rates, but are not commonly incorporated into models.

### Trait trade-offs

The presence of a vast diversity of phytoplankton species demonstrates that there is no one species that is a super competitor, dominating all others. This observation implies there is structure in the traits of species and has spurred a search for trade-offs among trait values and mechanistic explanations for these trade-offs. A well-developed understanding of the interrelationship of various trait values will help modelers constrain parameter values and produce better models. Many, but not all, of the identified trait trade-offs are related to cell size.

Large cells, particularly diatoms, appear to be especially adept at escaping grazing pressure and forming large blooms. It is hypothesized this is a consequence of size-specific grazing rates. Larger phytoplankton species are grazed by larger zooplankton species that have longer generation times compared to smaller predators. Thus large phytoplankton species have a growth advantage in certain habitats relative to smaller cells whose abundance is more tightly regulated by faster-growing grazers. A compilation of growth rate and nutrient acquisition traits from laboratory cultures showed species with higher maximum growth rates and maximum uptake rates have higher (worse) half-saturation constants and are poor competitors for that nutrient under low nutrient concentrations (have high *R*\* values). The correlation between high growth rate, high maximum nutrient concentration, poor half-saturation constants, and *R*\* is primarily due to the biophysical constraints associated with cell size (Fiksen et al. 2013; Litchman et al. 2007).

Some trade-offs are not primarily linked to cell size. In low nutrient environments, there is some evidence that the cell surface may limit the number of transporters, leading to a trade-off between the ability to acquire one nutrient compared to another (Edwards et al. 2011). Species that invest in mechanical protection against grazing by building armored cell walls (diatoms, coccolithophorids, dinoflagellates) often pay a metabolic cost for this armoring. The metabolic costs associated with producing armored covering are not well worked out. The increase in cell density due to calcification or silicification will mean that the rest of the cell must be less dense (or have other strategies) to maintain neutral buoyancy or their mean residence time in the upper mixed layer will be reduced and they will have increased rates of export out of surface waters.

### Trait differences across phytoplankton functional types

There are known and hypothesized differences in several traits across the major phytoplankton functional types. The binary traits (e.g., presence or absence of N_2_-fixation, silicification, calcification) define strong differences between functional types. The quantitative traits (e.g., maximum growth rate, nutrient affinity, average size) are thought to vary across functional types, but the amount of within-type variation in these trait values makes the differences much less certain. Although there may be meaningful differences in the quantitative traits across functional types, the available laboratory and field data indicate many of the quantitative traits overlap substantially across species from the different functional types.

Covariation among traits dominates the way functional types are commonly conceived. For example, many nitrogen-fixing cyanobacteria are relatively slow growing for their size and have high iron requirements due to the energy and iron requirements associated with nitrogen fixation. The common non-nitrogen fixing *Synechococcus* and *Prochlorococcus* have slow growth rates for their size (Raven et al. 2006), especially *Prochlorococcus*, and many *Prochlorococcus* strains are adapted to low irradiance and have lost the ability to use nitrate (Moore 2002; Zinser et al. 2007). Many diatom species under nutrient-sufficient conditions and adequate irradiance have fast maximum growth rates, high maximum nitrate uptake rates and lower half-saturation constants for nitrate than species of similar and even smaller size from other functional types (Litchman et al. 2007). Coccolithophores, often defined by the common species *Emiliania huxleyi*, can produce high levels of DMSP, have intermediate maximum growth rates, which are generally slower than diatoms, and a relatively low half-saturation constant for nitrate (Litchman et al. 2007). Differences in photosynthetic efficiency, maximum photosynthetic capacity, and ability to tolerate high irradiance are not well established across taxonomic groups. It has been hypothesized that the ability to tolerate high irradiance may shape the size structure of phytoplankton communities and that the high light tolerance of *Emiliania huxleyi* and *Phaeocystis* sp. (Loebl et al. 2010; Merico et al. 2004; Schoemann et al. 2005) and inability to tolerate high light by certain *Prochlorococcus* and *Synechococcus* stains may be key traits determining their temporal and spatial biogeographies (Six et al. 2007). There is also increasing evidence of functional type (especially size) differences in physiological response to different concentrations of dissolved inorganic carbon and pH but these traits are rarely explicitly considered in biogeochemical models (Wirtz 2011; Wu et al. 2014). Diatoms and coccolithophorids have higher sinking rates for their size relative to the other species due to their inorganic cell coverings. Many dinoflagellate species exhibit slow autotrophic growth rates and have high half-saturation constants for nitrate but are mixotrophic. There is relatively less work characterizing the traits of the Chlorophyta (or green algae) but there is some evidence they have better (smaller) half-saturation constants for ammonium than for nitrate (Litchman et al. 2007).

## Challenges using traits to represent functional types

### Challenges estimating average trait values for phytoplankton functional types

We know very little about the best ways to summarize a diverse group of species by a set of traits for a functional type. At present, modelers use a variety of approaches to scale up from traits of species to traits of functional types. Typically an average trait value is gathered from laboratory studies on a few key species from a functional type. A basic challenge in assigning trait values for phytoplankton functional types is that most phytoplankton species have not been studied in the laboratory and those species that have been studied document a great deal of variability in trait values within functional types.

There is extensive, order of magnitude variability in trait values across species within functional types under controlled laboratory conditions (Litchman et al. 2007). This variation is reflected in a large range of trait values used for individual phytoplankton functional types across different models (Table 3). For example, maximum growth rate values reported for nitrogen-fixing cyanobacteria range from 0.04 to 0.4 d^-1^, the half-saturation constant for phosphate for diatoms ranges from 0.00125 to 0.65 μmol L^-1^, and the half-saturation constant for iron for coccolithophores ranges from 0.02 to 2 nmol L^-1^. Differences in trait values across models arise because there is not enough laboratory data to confidently estimate trait values and because there is significant phenotypic plasticity and genetic variability in the trait values of interest across relevant species within a functional type. Variability in trait values among species and studies within the same functional type raises challenging questions about the appropriate trait values to use to represent functional types in models and how model traits should be defined and interpreted. The importance of variability in trait values within a functional type in biogeochemical models has not been sufficiently explored.

One solution to the problem of variability is simply to define trait values for a functional type by averaging trait values over all species with data within each type. This approach would work well for a functional type consisting of a few species all with similar trait values or if the species examined are similarly ecologically dominant. For diverse groups, such as the diatoms, or many groups with wide ranges of trait values, this approach is likely to lead to unrealistic trait values which do not represent the functional type well. Non-linear relationships among different traits can lead to an average for a suite of traits that is not representative of any species and is not characteristic of the functional type. Another challenge is that seasonal succession in a community may result in changes in functional traits for communities and perhaps for functional types (Weithoff and Gaedke 2016). A hierarchical Bayesian approach that quantified trait averages at species and higher taxonomic levels may provide better estimates of trait values from trait databases and enable a clear quantification of the uncertainty in our knowledge of trait values at various taxonomic levels (for an example see Finkel et al. 2016) and spatial and temporal scales. Another approach to finding trait values for functional types is to choose an exemplar species, which is either well studied or ecologically dominant, and simply use its trait values to represent the entire functional type (Table 3). *Emiliania huxylei*, for example, may be a stand-in for coccolithophores in general. Even this solution is only partial, because many ecotypes can be ecologically quite distinct within a species; for example, we know there are many different ecotypes of *E. huxleyi* (Iglesias-Rodriguez et al. 2006), *Prochlorococcus* and *Ostreococcus* (Kashtan et al. 2014; Rocap et al. 2003; Rodriguez et al. 2005).

### Challenges posed by acclimation and adaptation

Almost all biogeochemical models with phytoplankton functional types assume that traits are absolutely fixed. There are two problems with this assumption: phytoplankton can physiologically acclimate, within limits, to different environmental conditions, and phytoplankton can evolutionarily adapt to new conditions.

Many phytoplankton species have the capacity to alter their cellular physiology on time scales of seconds to days to improve their fitness under a range of environmental conditions. This means that traits, such as those characterizing nutrient acquisition, light absorption and photosynthesis, for individual species and functional types, may vary significantly under different environmental conditions. For example, there is evidence that nitrate uptake rates and photosynthetic parameters such as the chlorophyll-a specific light absorption coefficient, quantum yield of photosynthesis, maximum photosynthetic capacity, and *E_k_* vary with irradiance (Anning et al. 2000; Dugdale and Macisaac 1972; Falkowski and Laroche 1991; Lomas and Gilbert 1999). There may also be important differences across species and functional types in how fast traits track changes in environmental regimes. Traits are usually determined in the lab under equilibrium conditions, but natural communities are rarely at equilibrium, and environmental variability plays a crucial role in population dynamics. We may need to determine how nutrient and light acquisition and maximum growth rate change when cells are moved from replete to limiting conditions or vice versa (Grover 1991). One approach is to explicitly determine and add traits for acclimation. For example, Gregg et al. (2003) in a three-dimensional biogeochemical model allow the saturation irradiance, *E_k_*, to vary with irradiance. An alternate modeling approach, developed in the past decade, includes many species with similar but varied sets of trait values in each functional type and allows the environmental and biological conditions to empirically select which species, as defined by their trait values, are ecologically important (Follows et al. 2007). This permits different ecotypes to succeed in different parts of the ocean, even if locally much of the potential diversity is absent due to competitive exclusion. While it is possible to generate large numbers of species with trait values selected uniformly over some range, it would be helpful to have more knowledge about the kinetics and flexibility in key traits from lab studies and the true distribution of trait values in nature.

In addition to physiological acclimation to changes in the environment, phytoplankton may be able to adapt to changes in conditions on timescales of less than a decade. Short generation times and large population sizes mean that the evolutionary potential for phytoplankton is huge, and we have a poor understanding of the constraints on phytoplankton evolution. Experimental evolution in the lab has demonstrated that phytoplankton have the capacity to evolve when exposed to changes in CO_2_ concentration (Collins and Bell 2004; Lohbeck et al. 2012). The consequence of assuming traits are fixed when they can change through evolution is that predictions will likely over-estimate community change in response to changing climatic and environmental conditions (Irwin et al. 2015). Considerable effort will be required to identify and parameterize traits describing acclimation and adaptation that are most important to incorporate into models. Trait evolution is just beginning to be incorporated into large-scale biogeochemical functional type models using adaptive dynamic methods (Sauterey et al. 2014). There is hope that evolutionary experiments and an understanding of mechanistic reasons for trait value trade-offs will guide our forecasts of phytoplankton evolutionary potential and can eventually be incorporated into models, but there is much work to be done.

Several other open questions about trait modeling still need to be addressed. Traits may have very different meanings in two different models because of structural differences in the way the models are formulated. Models vary in their description of physiological responses to environmental conditions and interactions among factors, for example the way nutrient limitation affects growth. Some models use Liebig’s law of the minimum, others represent more complex co-limitation mechanisms, and some may have interactions between irradiance level and nutrient availability. This is a critical challenge for trait models and means that the idealized trait being a fundamental, fixed character of a species is not faithfully represented across models. New approaches such as Bayesian model selection may allow the community to resolve questions about the best way to model key processes (Anderson 2005). We have increasing evidence that species interactions, including mutualistic and allelopathic interactions, may be very important in structuring phytoplankton communities (Sher et al. 2011). Since our knowledge of interactions among traits is very incomplete, we are struggling to realize the full potential of phytoplankton trait models.

The promise of phytoplankton trait modeling is that knowledge of traits will allow us to project the biogeography of functional types and the consequences for biogeochemical cycles and ecosystem function. We have insufficient data on which traits are most important to project changes in phytoplankton functional type biomass, the appropriate values to use for these traits, and how trait values change with physiological acclimation and evolutionary adaptation. New analyses of field data that include species and environmental data may provide the guidance needed to identify key traits and species or communities that require additional study in the field and lab.

## Using field data to identify relevant traits and estimate trait values

### Why use field data?

There are vastly more species in natural communities than can be practically studied in the lab. Although we have a good idea of some of the important traits shaping community structure, we are not certain we have identified all the important core traits required to model a natural community. Natural communities present a much richer array of conditions than can be replicated in the lab. Analyses of field data could identify the species, traits, and environmental and biotic conditions that require more extensive evaluation in the lab. Methods are being developed so that traits can be determined from observations of natural communities, which may enable us to identify the most relevant traits and trait values. Further research is required to determine if traits derived from field data are as robust as the lab-derived traits and which are most useful for modeling. Instead of identifying individual species traits and developing trait models to ultimately obtain realized niches for species or functional types, it is possible to determine the realized niche of species and functional types directly from observations of natural populations (Box 1). Whether we estimate traits or niches, the dominant species and ecotypes that make up the natural communities are studied as opposed to model species easily cultured in the lab. Natural variability and biotic interactions are directly incorporated into the data analyzed.

#### Box 1 What is a niche? How are niches and traits connected?

Current definitions of a species’ niche describe it as the set of environmental conditions, or hypervolume, where the mean fitness of individuals of that species is greater than one. That is, the niche is the set of environmental conditions where the species can persist. This formulation is due to Hutchinson (1957; 1978) and differs considerably from earlier concepts developed by Gause, Elton, and Grinnell. Before Hutchinson, the niche was not a purely abstract set of environmental conditions, but was tied to a physical location, so that two coexisting species could literally occupy the same niche. Hutchinson identified the physical space occupied by a species as its biotope. The mapping of niches onto physical spaces (the biotope) and the inverse problem of identifying niches from biotopes is the central goal of species distribution modeling (Colwell and Rangel 2009).

Hutchinson distinguished the fundamental from the realized niche. The fundamental niche is the set of conditions in which a species can persist without considering interactions with other species. For phytoplankton, this is the niche commonly measured in the lab from uni-algal cultures. The realized niche incorporates the effects of interactions with other species. For example, two species with overlapping fundamental niches may differ in their competitive ability where niches overlap, and one species may exclude the other from some of its niche, or the two species may coexist. Facilitation between species may even expand the realized niches beyond the fundamental niches of either species (Bruno et al. 2003). The realized niche can also be reduced relative to the fundamental niche because of dispersal limitation or the absence from the environment of some part of the fundamental niche. This last reason presents a particular challenge since future oceans are likely to contain combinations of environmental conditions present nowhere in today’s ocean; some of these conditions may be part of a species’ fundamental niche, but can’t be part of any realized niche measured today.

Mechanistic models based on traits can be used to predict the niche and biotope of a species, but the model must describe the fitness of individuals and how they interact with their environment and other organisms sharing the same habitat (Kearney 2006; Kearney et al. 2010). As we explain in the main text, this plan may be difficult for individual species of phytoplankton, because we lack considerable information both about species’ traits and species interactions. While the trait-based modeling approach is very attractive, it is not clear whether fundamental niches and traits from lab studies or empirical niches from species distribution models will be most useful. It is probably best to see them as complementary. When more phytoplankton trait values are known, it may be possible to adapt a hybrid “trait niche” approach now being explored for plants (Violle and Jiang 2009).

### How can we identify traits and niches of phytoplankton functional types from field data?

Many methods are available to characterize the traits and niches of species and functional types in natural populations. Two broad categories are parameter estimation from mechanistic models and species distribution models (SDMs). Time-series data with sufficient temporal resolution can be used in combination with mechanistic models to estimate traits governing growth and loss terms, but these data are rare. Mechanistic models are usually differential equations that describe the rate of change of population abundance or biomass. These models are commonly formulated using functions for resource acquisition, growth, and loss rates (Fig. 1), ranging from very simple phenomenological parameterizations to descriptions of intricate, multistep mechanistic processes. Key parameters in these models are the phytoplankton traits, which can either be estimated one at a time from controlled lab experiments or jointly using an optimization approach. Mechanistic models have been used to estimate daily growth and loss rates for *Synechococcus* off the coast of New Jersey at LEO-15 (Sosik et al. 2003) and for three groups of picophytoplankton in the central equatorial Pacific (André et al. 1999). The frequency of observation will determine key aspects of even apparently easily interpreted traits such as growth rate. Sosik et al. (2003) estimated population counts and sizes several times a day,so the estimated growth rates closely approximate the true growth rate of *Synechococcus* at this site. We have experimented with obtaining trait values from weekly data but because of the greater time between samples, these growth rates will incorporate many loss terms and will be considerably smaller than the maximum growth rate of the species present in the community (Mutshinda et al. 2016). It is probably impossible to obtain meaningful growth rates from monthly time-series data since phytoplankton generation times are much less than one month.

Species distribution models (SDMs) or bio-climate envelope models can be based on statistical tools such as generalized linear models (GLMs (Guisan et al. 1999)), non-parametric statistical approaches such as generalized additive models (GAMs, e.g., (Xiao et al. 2016)), or more complex machine learning methods such as random forests (Evans et al. 2011) or maximum entropy methods (Elith et al. 2011). They can use time-series data or any coincident species and environment data. The BioMod approach combines many of these types of methods in an effort to test for robustness of the SDMs (Thuiller et al. 2013; Thuiller et al. 2009). These methods are generally non-mechanistic, although the predictor variables incorporated into the models can be clearly linked to mechanisms. In general, SDMs are designed for prediction of species or functional type presence or abundance rather than for estimation of traits but the prediction can be used to generate a description of the niche (see Box 1). In this way SDMs provide the end product of what trait-based models are designed to produce: an estimation of how environmental conditions affect presence and abundance. Below we provide examples of how phytoplankton functional type niches can be derived from field data and provide new insight into niche differentiation across functional types and the evolutionary capacity of phytoplankton niches to adapt to climate forcing on decadal scales.

We have used a maximum entropy approach (MaxEnt) to develop SDMs and extract multivariate niches for 119 species of phytoplankton (diatoms and dinoflagellates) from 60 years of data collected by the North Atlantic Continuous Plankton Recorder program (CPR) conducted by the Sir Alister Hardy Foundation for Ocean Sciences (Irwin et al. 2012) and 67 species from Station Cariaco over 15 years (Irwin et al. 2015). MaxEnt is a machine-learning statistical method that can be used to estimate the probability that a species is present under a given set of environmental conditions (Elith et al. 2011; Phillips and Dudík 2008). MaxEnt is notable because it operates on species occurrence data, not abundance or biomass data, and it is designed to not require knowledge of conditions under which the species is absent. This is particularly important for time-series such as the phytoplankton presence and absence data in the CPR dataset because biomass loading on the sampling silk of the CPR sampler is variable and known to affect detectability of some species. The probability of a species is present as a function of environmental conditions can be condensed into summary statistics such as the mean of the niche (Irwin et al. 2012, Box 2). A similar analysis has been done on the MAREDAT database that spans more taxonomic groups and a larger geographic region (Brun et al. 2015; Buitenhuis et al. 2013). We found diatoms, compared to dinoflagellates, are more likely to be found in colder, more nutrient rich water with lower light levels and deeper mixed layers both in the North Atlantic and at the CARIACO time series station in the Caribbean (Irwin et al. 2012; Mutshinda et al. 2013, Fig. 2). The functional-type level differences in niche appear to be robust across regions and investigators (Brun et al. 2015; Irwin et al. 2015; Mutshinda et al. 2013). These analyses indicate that there are many distinct niches for phytoplankton, but that species belonging to the same phylum or functional type tend to have more similar niches than species from different phyla or functional types. A number of species within each of the functional types have niches more similar to other functional types, and within a functional type niches of individual species do often overlap, but there are many species with narrow niches quite distinct from other species. Once the realized or fundamental environmental niches from phytoplankton species have been identified they can be used to project how species and functional type biogeography may change with climate over the next decades to centuries. Our projections indicate, for example, that climate change over the next century will stimulate many species to migrate northeast in the North Atlantic (Barton et al. 2015).

#### Box 2 Cartoon of phytoplankton niches in three dimensions

The niches of four hypothetical species (ellipsoids) for two different functional types (dark and light shading) are shown as a function of three environmental conditions (temperature, irradiance, and nitrate concentration; arbitrary scales for each). Species frequently exhibit considerable overlap in niche space and a key element of the functional type approach is that niches of species from the same functional type are more similar to each other, on average, than species from different functional types.

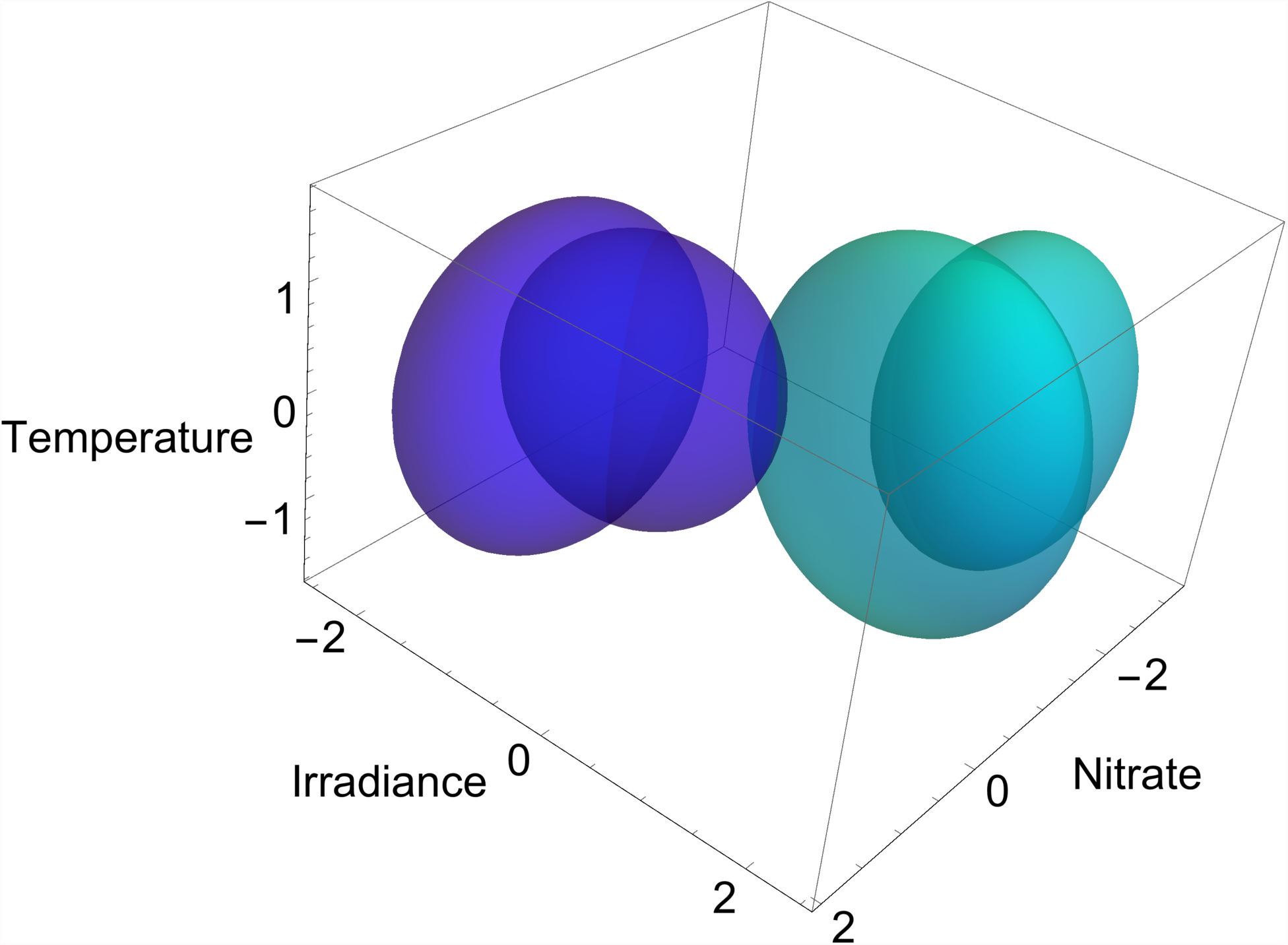

**Figure 2.**
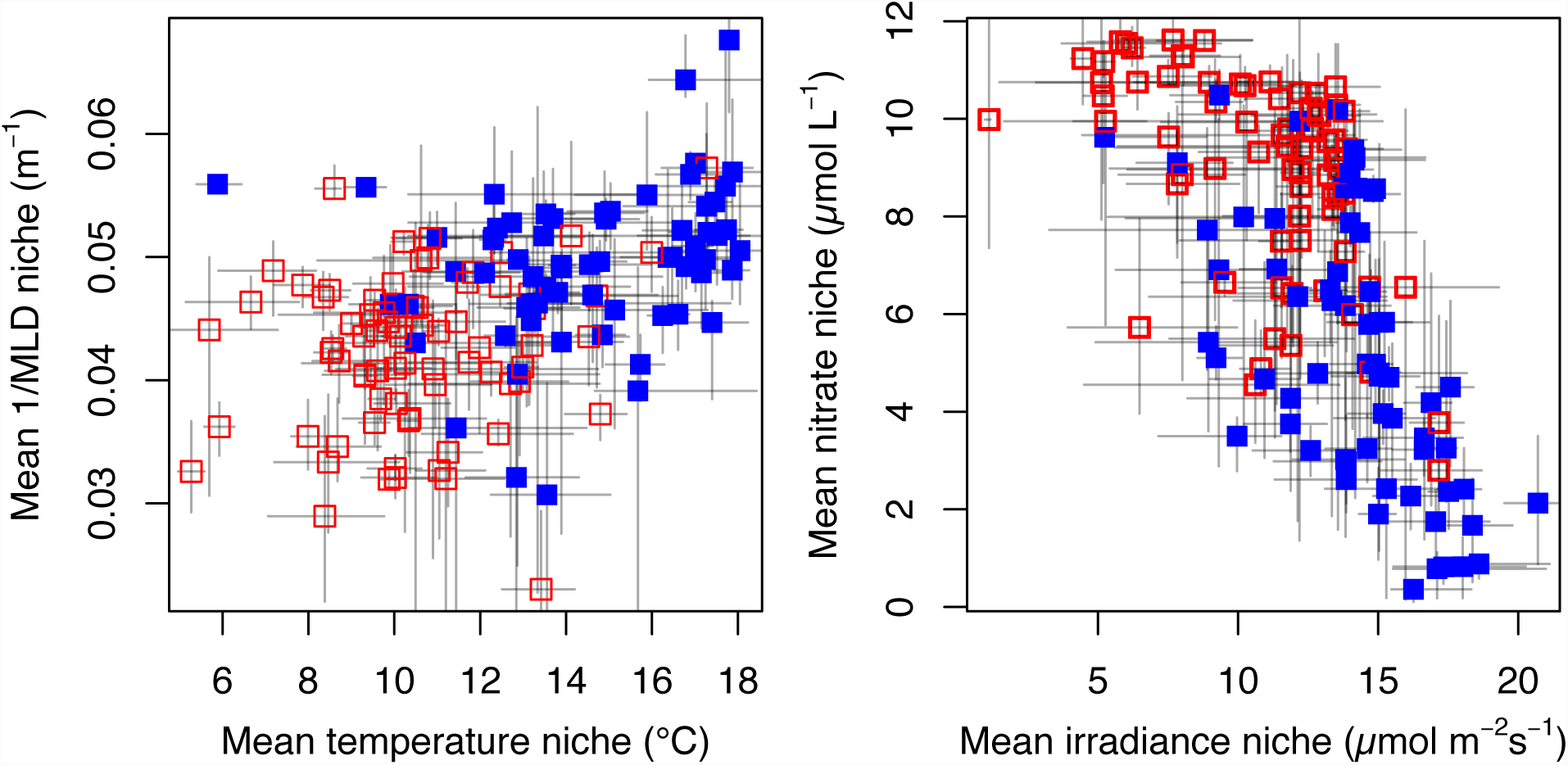
Mean niches estimated from time-series data. Mean niches derived from MaxEnt species distribution models for 69 diatom species and 50 dinoflagellate species from the North Atlantic Continuous Plankton Recorder. We summarize the niche for each species by its mean. Left panel: the reciprocal of mixed layer depths versus temperature. Right panel: nitrate concentration versus mean irradiance in the mixed layer. The traits separate the two functional types, diatoms (open squares, light color) and dinoflagellates (filled squares, darker color), although there is a great deal of variation within the functional types. Error bars are 95% confidence intervals on the means estimated by bootstrap resampling. For full details see Irwin et al. (2012).

### Are phytoplankton niches stable over time?

Statistical analysis of natural populations yields realized niches that can be quite different from the fundamental niches of the same species as measured in the lab, one species at a time. These differences can be due to interactions between species, the absence of part of the fundamental niche from the habitat studied, or even dispersal limitation. Any of these factors could change over time or as the environment changed, so it is reasonable to ask if realized niches are stable as the environment changes. In addition, since phytoplankton have large population sizes, standing-stock diversity, and short generation times, they have a large evolutionary potential, and changes in the realized niches could be due to evolution. Station Cariaco, in the Caribbean Sea is an ideal location to test the temporal stability of realized niches for phytoplankton. Unlike the North Atlantic CPR for which we used climatological environmental data, the Station Cariaco time-series has simultaneous monthly observations of temperature, salinity, and macronutrient concentrations. These data document an increase in the mean upper mixed layer temperature of about 1°C, a small increase in mean irradiance in the mixed layer, and a pronounced increase in the number of months with very low nitrate concentration in the sea surface over the past 15 years (Taylor et al. 2012). We have used MaxEnt and hierarchical Bayesian models to identify the niches of the dominant phytoplankton species at Station Cariaco (Irwin et al. 2015; Mutshinda et al. 2013) and to test if the realized niches of phytoplankton species have been constant or if they have changed over the last 15 years. Over a decadal scale, temperature and irradiance niches tracked changes in the environment, but for most species, the nitrate niches remained fixed despite a marked change in mean nitrate conditions (Irwin et al. 2015). Phytoplankton realized niches should not be assumed to be stable, and evolutionary changes in phytoplankton functional traits should be considered in climate and biogeochemical projections that extend more than a decade into the future.

## Should we model functional types or individual species?

Most studies incorporating phytoplankton functional types simply assume that species can be naturally grouped into functional types. Few studies have explicitly tested how well functional types represent individual species within the functional grouping. Neutral theory can be used to test if individual species dynamics are consistent with the average functional type dynamics. In recent years a neutral theory of biodiversity and community dynamics has been explored in stark contrast to niche models (Hubbell 2001). The neutral theory hypothesizes that community structure is determined by dispersal and random fluctuations in abundance rather than a filtering of species by environmental conditions. If the biomass dynamics of a species is a random walk within the functional type this indicates the species behaves like the average of the functional type (See Box 3). If the biomass dynamics of a species is not a random walk, we conclude that the species does not vary neutrally within its functional type.

### Box 3 Random walks and neutral theory

The idea of a random walk originated from observations of Brownian motion: particles moving in many different directions as a result of a vast number of random collisions. Mathematically a random walk along one dimension is a sequence of random variables representing the position of a particle that moves one step in either direction at each time step. The random variables are memoryless (the defining characteristic of a Markov process), so that the direction of movement does not depend on the past history of the process. Four sample random walk trajectories starting at the same location are illustrated.

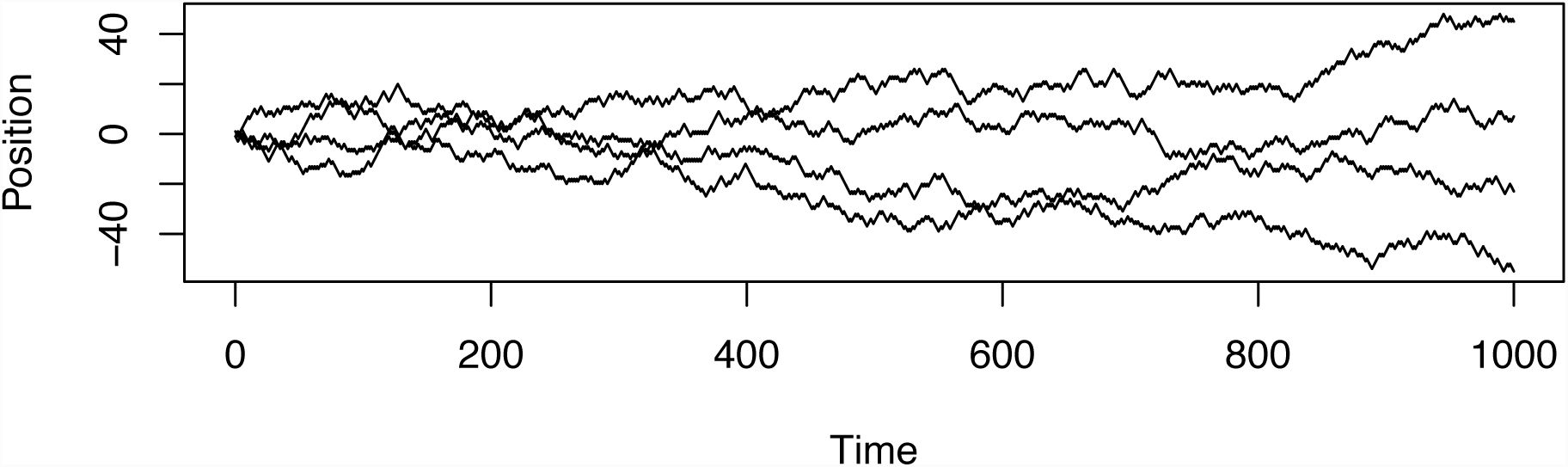

A random walk can be used as a model of species population dynamics consistent with the neutral theory of ecology. Species are neutral if their populations fluctuate without regard to their trait values or the environmental conditions, and this leads to species abundance following a random walk. For phytoplankton, we know that certain environmental conditions favor one functional type over another, for example, diatoms over dinoflagellates. So the community biomass dynamics of functional types are not neutral. Individual species may still have neutral biomass dynamics within its functional type, for example many diatom species may exhibit random walk behavior relative to the total biomass of diatoms. We refer to this restricted example of neutrality as species neutrality relative to the biomass envelope of its functional type.

We used seven years of weekly data on 57 diatom species and 17 dinoflagellate species with coincident environmental data from the L4 time series from the Western Channel Observatory (Plymouth, UK) to evaluate if a simple trait based model could predict week-to week changes in the aggregate biomass in diatoms and dinoflagellates and if the biomass dynamics of individual diatom and dinoflagellate species were neutral, i.e., random changes within their functional type. We found a simple set of functional traits (maximum growth rate modified by irradiance, temperature and a density dependent loss rate), with distinct values for the diatoms and the dinoflagellates, describes the vast majority (96%) of the temporal variation in biomass for those two functional types (Mutshinda et al. 2016). On average the diatoms had higher biomass accumulation under lower temperatures and irradiances than the dinoflagellates at the L4 site.

We found that a few species had strongly neutral dynamics, but for most species, their dynamics were neutral within their functional type biomass envelope about half the time. The neutral theory has also been tested for phytoplankton along the Atlantic Meridional Transect (AMT). Phytoplankton communities along the AMT also appear to be partially determined by environmental filtering and partially by neutral dynamics at the functional type level (Chust et al. 2013), although the explanatory power of both mechanisms was limited in this case. Taken together these results mean that while there are differences in traits from species to species within phytoplankton functional types, most species behave neutrally within their functional type most of the time. In other words the majority of species within their functional types are well represented by average trait values estimated from aggregate biomass, although variability at the species level means that biomass of individual species may be difficult to predict.

These results indicate that phytoplankton species naturally cluster into taxonomically defined phytoplankton functional types due to a similarity in traits arising from a shared phylogenetic history. It is easier to predict changes in abundance of phytoplankton functional types than the biomass of individual species within the functional types. These results suggest that for broad-scale ecological questions it may be best to model phytoplankton communities at the functional-type level than at the species level because the extra trait values required to develop a species model are not needed to obtain predictions about functional types. Furthermore this approach presents a straightforward way to estimate the trait values of a functional type directly from community-level data in natural populations (Mutshinda et al. 2016), rather than the dubious practices of selecting a single representative species or averaging traits from several representative species.

## A way forward

Functional types are used to gather many phytoplankton species into a small number of categories and to focus attention on functional-type level differences in physiology, biogeography, ecological function, and biogeochemical roles. Analyses and projections using phytoplankton functional types are generally more informative and more robust than approaches that use species. Functional types provide the joint benefits of better predictions, simpler data requirements, and reduced complexity.

The reasons for the relative success of the functional type approach compared to individual species models for predicting biogeography, productivity, and biogeochemical cycles are not completely clear. Phytoplankton communities have complex dynamics, and we often lack information about the physical and chemical data affecting the maximum and realized growth rate and about the grazer community and other factors determining loss rates. Averaging over species in a functional type may remove the need for some of the detailed data we lack and may be sufficient for many relevant ecological and biogeochemical scale questions. The dominant species within a functional type may be similar enough to each other that it is not necessary to resolve the species-level differences for some questions.

To make functional type modeling possible, researchers attempt to determine appropriate trait values for a whole functional type. Traits with binary or categorical values (e.g., presence or absence of silicification, calcification, N_2_-fixation) may be the most suited to defining functional types as they are more highly conserved within evolutionary lineages than many of the quantitative traits. Average trait values for quantitative traits (e.g., maximum growth rate, half-saturation constants) are different, in some average sense, across some functional types, but the values of many traits vary by more than 10-fold across species within a functional type. Some models make realistic large-scale predictions of major phytoplankton functional types, but the vastness of the ocean means that they are difficult to validate with sufficient data. Models differ in the functional types, the traits, and the trait values. Trait values used in these models are often tuned to available observations, which may account for similarities in predictions despite the variation in trait values used across models. Structural differences exist among models, which further complicates comparisons of models and trait values with each other and with trait values derived from lab experiments. Since the quantitative trait values are crucial parameters in phytoplankton functional type models, we need a better understanding of trait variation and co-variation within and among functional types. Much more work could be done to identify the fundamental traits most suitable for specific research questions and to explore the consequences of different model formulations and the sensitivity of results to model formulation and trait values.

We have identified several challenges for researchers seeking to use phytoplankton functional types and functional traits in models. Differences between trait values in the lab compared to values used in models and variation in values used in models suggests that we are still struggling to find a suitable representation of traits and trait values for functional types in models. Modeling efforts have concentrated on incorporating trait values we know the most about, but we have not spent enough effort determining which traits are crucial for ecological and biogeochemical modeling. Sensitivity analyses systematically and quantitatively explore the consequences of varying model formulations and parameter values and can be used to assess the consequences for predictions of our uncertainty for specific trait values.

Each research question has its own requirements for spatial and temporal scales and these influence how predictive models should be formulated and perhaps which traits will be most relevant. Depending on the time-scale, physiological acclimation or evolutionary adaptation may be very important, but we know very little about how to best incorporate these processes into models. In highly variable natural environments, physiological time-scales of acclimation may be important for projecting community dynamics over days to weeks, but little work has been done to connect physiological acclimation studied in the lab to models describing natural populations. Lab experiments can demonstrate the capacity for phytoplankton species to adapt evolutionarily to changes in their environment and this capacity needs to be connected to long-term field data to determine the evolutionary potential of natural populations. Each of these challenges is an opportunity for new insights and creative collaborations.

There is still much to be learned about the niches and traits of phytoplankton species and functional types from analyses of field data. New analyses of field data may be able to identify the key minimum set of traits and key species within functional types that merit further intensive laboratory, mesocosm, and modeling work. Comparisons of trait values from time-series studies with trait values determined in the lab on species isolated from these time series could provide valuable insight into the relationship between the fundamental and realized niches of ecologically dominant phytoplankton species and how to most appropriately assign trait values to phytoplankton functional types. Mesocosms incorporating these species could be integrated with modeling efforts to test model formulations and assess the importance of individual traits and trait variation to the success of model outputs. A tighter integration of statistical analyses of field data, models, and lab studies will certainly improve our definition of phytoplankton functional types and the value of phytoplankton functional type modeling.

## Summary

1. Phytoplankton functional types are aggregations of species into groups that perform broadly similar ecological or biogeochemical functions.
2. Trait values for individual phytoplankton species measured in lab may not be generally representative of the trait values for the corresponding functional type in field conditions and may vary over time and region.
3. We have insufficient data to reliably quantify trait values for the wide diversity of phytoplankton species.
4. A possible solution is to directly estimate trait values for phytoplankton functional types from field data using statistical modeling tools.
5. Evidence suggests that ecological modeling of phytoplankton functional types may be more successful than efforts to model individual species. Trait modeling of functional types rather than species is likely to be the best way forward.

